# A performance assessment of relatedness inference methods using genome-wide data from thousands of relatives

**DOI:** 10.1101/106013

**Authors:** Monica D. Ramstetter, Thomas D. Dyer, Donna M. Lehman, Joanne E. Curran, Ravindranath Duggirala, John Blangero, Jason G. Mezey, Amy L. Williams

**Author notes:** Correspondence (M.D.R.) or (A.L.W.).

## Abstract

Inferring relatedness from genomic data is an essential component of genetic association studies, population genetics, forensics, and genealogy. While numerous methods exist for inferring relatedness, thorough evaluation of these approaches in real data has been lacking. Here, we report an assessment of 12 state-of-the-art pairwise relatedness inference methods using a dataset with 2,485 individuals contained in several large pedigrees that span up to six generations. We find that all methods have high accuracy (~92% – 99%) when detecting first and second degree relationships, but their accuracy dwindles to less than 43% for seventh degree relationships. However, most IBD segment-based methods inferred seventh degree relatives correct to within one relatedness degree for more than 76% of relative pairs. Overall, the most accurate methods are ERSA and approaches that compute total IBD sharing using the output from GERMLINE and Refined IBD to infer relatedness. Combining information from the most accurate methods provides little accuracy improvement, indicating that novel approaches—such as new methods that leverage relatedness signals from multiple samples—are needed to achieve a sizeable jump in performance.

---

The recent explosive growth in sample sizes of genetic studies has led to an increasing proportion of individuals with at least one close relative in a dataset, necessitating relatedness detection. As the number of pairs in a sample grows quadratically in its size, for a constant rate of relatedness among pairs, proportionately more individuals will have close relatives in large datasets. This pervasiveness has relevance to nearly every genetic analysis performed in moderate to large scale data, including trait mapping and population genetics. In particular, inferring relatedness between samples^1–3^ is essential to avoid spurious signals in genetic association studies^4–6^; empowers linkage analysis by enabling the correct specification of pedigree structures^7–9^; facilitates identification of relatives in the context of forensic genetics^1,10,11^; and is needed to account for or remove relatives in population genetic analyses^12–14^. Relatedness estimation has also drawn the interest of the general public via companies that offer genetic testing services and advertise their ability to find customers’ relatives, thus allowing individuals to explore their ancestry and genealogy. The broad utility of relatedness detection has motivated the development of numerous methods for such inference. These methods work by estimating the proportion of the genome shared identical by descent (IBD) between individuals^1,3^ or a closely-related quantity, where an allele in two or more individuals’ genomes is said to be IBD if those individuals inherit it from a recent common ancestor^2^. Characterizing the true relatedness of two or more samples is challenging for several reasons, including chance sharing of alleles between individuals who are only distantly related, and the fact that the distributions of IBD proportions for different relatedness classes overlap^2,15^ (e.g., first cousins and half-first cousins).

Motivated by the substantial need to identify relatives in modern samples, we present an evaluation of 12 state-of-the-art pairwise relatedness methods, each capable of scaling to analyze thousands of individuals, including seven that directly infer genome-wide relatedness measures^16–22^ and five IBD segment detection methods^23–27^ that we utilized to infer these quantities. To assess these methods, we used SNP array genotypes from Mexican American individuals contained in large pedigrees from the San Antonio Mexican American Family Studies (SAMAFS)^28–30^. Our analysis sample included 2,485 individuals genotyped at 521,184 SNPs (Supplemental Note) within pedigrees that span up to six generations, and with genotype data from as many as five generations of individuals. Given this large sample, including 13 pedigrees with >50 individuals (Supplemental Figure 1), numerous relatives exist, and we used these to evaluate the inference methods. In particular, we analyzed >3,700 pairs of individuals within each of the first through fifth degree relatedness classes, 816 and 73 sixth and seventh degree relatives, respectively, and more than three million pairs of individuals that are reported as unrelated (Table 1). Prior evaluations of relatedness inference methods included only a subset of the methods we evaluate, and either considered simulated data^17,18,20–22^ (which may not fully capture the complexities of real data), used small sample sizes^17,18,22,31^, or did not consider sixth and seventh degree relatives^17,18,20,22^. This analysis of real data from large numbers of up to sixth degree relatives, as well as dozens of seventh degree relative pairs, provides a comprehensive evaluation of existing pairwise relatedness inference methods.

**Table 1:** Number of pairs of individuals in the SAMAFS dataset that passed sample filters (Supplemental Note) and are reported to have relatedness between first and seventh degree or as unrelated. We combined reported monozygotic (MZ) twins with the set of first degree relatives.

| Degree | Number of Pairs |
| --- | --- |
| 1 | 4,969 |
| 2 | 6,625 |
| 3 | 8,241 |
| 4 | 7,636 |
| 5 | 3,794 |
| 6 | 816 |
| 7 | 73 |
| Unrelated | 3,051,598 |
| Total | 3,083,752 |

The performance metric for this study is the rate at which each method infers the pairs of samples to have the same degree of relatedness as that reported in the SAMAFS pedigrees. These reported relationships are generally reliable, and we filtered out relative pairs whose degree of relatedness is potentially inflated due to apparent relatedness on ancestral lineages reported as unrelated (Supplemental Note). Some programs directly infer the degree of relatedness^19^, while others infer a kinship coefficient ^17,18,20^ or a coefficient of relatedness^16,22^ (which is two times the kinship coefficient^32^), and the remainder instead detect IBD segments^23–27^ (Table 2). To infer the degree of relatedness from an estimated kinship coefficient, we use the mapping recommended in the KING paper ^17^ (Supplemental Table 1), which are ranges that use differences in powers of two for the relatedness degree intervals and are generally consistent with simulations 17.

**Table 2:** Properties of the 12 relationship inference methods we analyzed. Type indicates the inference methodology the program uses. Runtime is wall clock time to run the program with any additional time to run programs needed for input as indicated. We ran parallelized programs using the numbers of cores indicated in parentheses: total compute time for the parallelized programs is the runtime multiplied by the number of cores used. Input required from outside program indicates extraneous information needed to run the program. Programs that use either principal components, sample ancestral population proportions, or that use a model designed for multiple populations are indicated as accounting for population structure. “Y” indicates yes, “N” indicates no, and “NA” indicates not applicable. Runtimes are from a machine with four AMD Opteron 6176 2.30 GHz processors (64 cores total) and 256 GB memory. *Additional time to phase the data using Beagle 4.1 and run GERMLINE. ^†^Additional time to phase the data using Beagle 4.1. ^‡^.Additional time to obtain KING relatedness estimates; base PC-Relate time is the sum of time to run this method and PC-AiR^34^. ^§^Additional time to obtain ancestral population proportions using ADMIXTURE^35^.

| Method | Version | Citation Number | Type | Output | Parallelized? | Runtime (×cores used if $>1$) | Requires independent markers | Input required from outside program | Accounts for population structure |
| --- | --- | --- | --- | --- | --- | --- | --- | --- | --- |
| **ERSA** | 2.0 | 19 | IBD segment-based | Degree of relatedness | N | 14.3h + 96.3h ($\times 16$)* | N | IBD segments | NA |
| **fastIBD** | Beagle 3.3.2 | 24 | IBD segment-finding | IBD segments | N | 55.2h | N | NA | NA |
| **GERMLINE** (-haploid) | 1.5.1 | 23 | IBD segment-finding (Distinguishes IBD1 and IBD2) | IBD segments | N | 19.2m + 96.0h ($\times 16$)<sup>†</sup> | N | Phased genotypes | NA |
| **HaploScore** | NA | 27 | IBD segment-based | IBD segments | N | 2.4h + 96.3h ($\times 16$)* | N | IBD segments; phased genotypes | NA |
| **IBDseq** | r1206 | 25 | IBD segment-finding | IBD segments | Y | 33.1h ($\times 16$) | N | NA | NA |
| **KING** (KING-robust) | 1.4 | 17 | Allele frequency-based IBD estimate | IBD 0,1,2 proportions | N | 4.6m | Y | NA | Y |
| **PC-Relate** | 2.0.1 | 22 | Allele frequency-based IBD estimate | IBD 0,1,2 proportions | N | 8.9h + 4.6m<sup>‡</sup> | Y | Pairwise kinship coefficients | Y |
| **PLINK 1.9** | 1.90b2k | 16 | Allele frequency-based IBD estimate | IBD 0,1,2 proportions | N | 18.1s | Y | NA | N |
| **PREST-plus** | 4.1 | 33 | Allele frequency-based; uses linkage model | IBD 0,1,2 proportions | N | 178.9h | N | NA | N |
| **REAP** | 1.2 | 18 | Allele frequency-based IBD estimate | IBD 0,1,2 proportions | N | 3.8h + 2.8h<sup>§</sup> | Y | Ancestral population allele frequencies; sample ancestry proportions | Y |
| **Refined IBD** | Beagle 4.1 | 26 | IBD segment-finding (Distinguishes IBD1 and IBD2) | IBD segments | Y | 96.0h ($\times 16$) | N | NA | NA |
| **RelateAdmix** | 0.1 | 20 | Allele frequency-based IBD estimate | IBD 0,1,2 proportions | Y | 15.8h ($\times 16$) + 2.8h<sup>§</sup> | Y | Ancestral population allele frequencies; sample ancestry proportions | Y |

For IBD detection methods that report the number of IBD segments shared at a locus^23,26^—denoted IBD0, IBD1, and IBD2 for the corresponding number of copies that are IBD—it is straightforward to calculate a kinship coefficient^2^. This coefficient, *ϕ_ij_*, between a pair of samples *i, j* denotes the probability that a randomly selected allele in individual *i* is IBD with a randomly selected allele from the same genomic position in *j*. Let 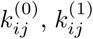, and 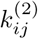 denote the proportion of their genomes that individuals *i*, *j* share IBD0, IBD1, and IBD2 respectively; then the kinship coefficient is 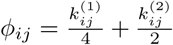. The proportions 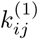 and 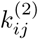 are simply the sum of the genetic lengths of the IBD1 and IBD2 segments, respectively, between samples *i, j* divided by the total genetic length of the genome analyzed. For the IBD detection methods^24,25,27^ that do not distinguish between regions that are IBD1 from IBD2, the proportion of the genome that is inferred to be IBD0 provides an alternate means of estimating the degree of relatedness (Supplemental Table 1), with the ranges of values here again from the KING paper ^17^. We classified pairs of individuals with lower kinship coefficients or higher IBD0 rates than indicated for the eighth degree range as unrelated.

The results from the analysis are shown in Figure 1, which depicts the proportion of sample pairs inferred to be within each of the degree classes that we considered (first through eighth degree and unrelated), separated according to their reported relatedness degree. All methods perform well when inferring first and second degree relatives, with accuracies ranging from 98.8% to 99.5% for first degree relatives, and from 92.8% to 98.6% for second degree relatives. However, the methods’ accuracies diverge for more distant relatedness, with the IBD segment-based methods generally having higher accuracy than those that rely on allele frequencies of independent markers. For example, for sixth and seventh degree relatives, the top performing IBD segment-based method has 58.1% and 42.5% accuracy, respectively, while the highest performing allele frequency-based method has an accuracy of only 44.6% and 27.4%, respectively. This general pattern applies to fourth and fifth degree relatives as well, although with less discrepancy between these two inference approaches for these closer relatives. The decreased inference accuracy of all methods for higher relatedness degrees is likely due to the exponential drop in mean pairwise IBD shared and an increased coefficient of variation for more distant relationships ^15,36,37^.

**Figure 1:**
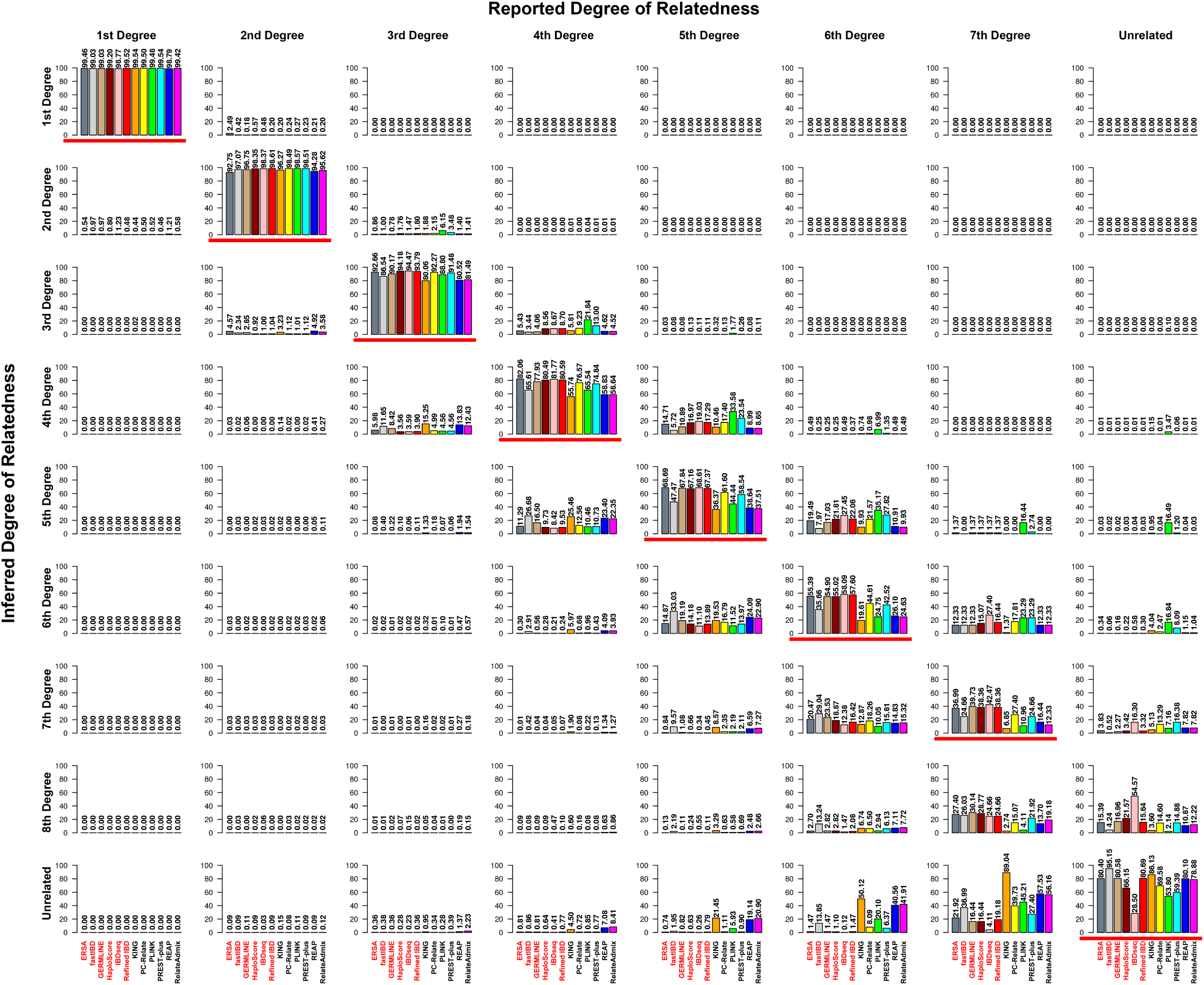
Performance comparison of the evaluated methods using the SAMAFS dataset. Bar plots indicate the percentage of sample pairs that are reported to have a given degree of relatedness and that are inferred to be related as the indicated degree. The bar plots are separated on the horizontal axis by the reported relatedness degree and on the vertical axis by inferred relatedness degree. For clarity, the plots list above each bar the inferred percentage that the corresponding bar depicts. Program names listed in red are IBD segment-based methods while those in black utilize allele frequencies for inference. Red horizontal bars under a bar plot indicate that the corresponding inferences agree with the reported relationships.

While the accuracies for exact inference of distant relatives are fairly low among all methods, the IBD segment-based methods (excluding fastIBD) are correct to within one degree of the reported relationship at a rate of ≥95.3% for sixth degree relatives and ≥76.7% for seventh degree relatives. At the same time, ERSA, GERMLINE, and Refined IBD classify ≥80.4% pairs of unrelated individuals correctly, and several other methods also correctly infer ~80% pairs of unrelated individuals, although many of these methods perform poorly when classifying reported relatives. The inference of ~20% of the more than three million unrelated samples as eighth degree or closer relatives suggests the presence of a non-trivial fraction of unreported relationships in these data. Alternatively, and perhaps more likely, many of these may be false positive relationships, as distinguishing pairs of unrelated individuals from fairly distant relatives is difficult. With the lower bound for eighth degree relatives being a total of 19.5 cM of IBD segments shared between individuals, spurious inferences at this level are possible, with IBD segments detected in regions subject to historical selection^38^ or with low SNP density potentially leading to inflated IBD proportions. In that regard, we note that some analyses of IBD reweight segments that overlap regions with excess IBD sharing in order to improve the reliability of overall sharing rates ^39,40^. Additionally, analyses that consider relatedness among the parents and/or children of inferred distant relatives have the potential to avoid some of these issues, and indeed, the recently developed relatedness classification method PADRE does analyze familial relatedness signals and shows improved accuracy^41^.

Overall, the most accurate programs for first through seventh degree and unrelated classification are ERSA, GERMLINE, and Refined IBD—all IBD segment-based methods. The improved accuracy of these methods may be due to their focus on identifying long stretches of identical haplotype segments that more readily discriminate recent shared relatedness from chance sharing of alleles. The IBDseq method, while performing well for inferring first through seventh degree relatives, infers a much larger fraction of pairs of individuals as related that are reported as unrelated, suggesting it may be biased towards detecting higher levels of IBD sharing than the other methods.

Noting that the SAMAFS consist of admixed Mexican American individuals, we examined the accuracy results among the allele frequency-based methods, several of which account for population structure. While IBD segment-based methods generally have the best performance and do not directly account for population structure, inferring IBD segments is computationally demanding, and considering the performance of more efficient allele frequency-based methods is of interest. Among all these methods, PC-Relate has the highest accuracy across all levels of relatedness, and it accounts for population structure using principal components (PCs) inferred from a set of samples with low relatedness ^22^. However, PREST-plus has only slightly lower performance than PC-Relate even though it does not account for population structure. PREST-plus implements a hidden Markov model (HMM) that enables it to leverage linkage signals to identify regions that are likely to be IBD between samples^21^. Therefore, although PREST-plus does not explicitly detect IBD segments, it leverages similar signals to the IBD segment-based approaches, which might enable it to be less susceptible to biases caused by ignoring the effects of population structure. Relatedness estimation that ignores population structure in admixed samples can produce either a positive or negative bias^22^. Consistent with this, PLINK infers many sample pairs to be more related than they are reported to be, and, at the same time, infers substantial fractions of fourth through seventh degree pairs as unrelated. KING also dramatically underestimates relatedness, presumably because it assumes that all samples derive from one of several homogeneous populations—a model that is inappropriate for recently admixed samples^17^. We also examined results from the version of KING that assumes a single homogeneous population and its accuracy profile more closely resembles that of PLINK (not shown).

Because the relatedness within SAMAFS has the potential to confound methods that characterize population structure ^34^, we further analyzed the performance of several methods using a dataset consisting of the SAMAFS samples together with a diverse set of HapMap individuals ^42^ (Supplemental Note; Supplemental Figure 4). This combined dataset yields inferences of sample ancestry proportions that are strongly correlated with those inferred in a reduced dataset that has only low level relatedness (Supplemental Note). Using this sample, the accuracies of both REAP and RelateAdmix improve significantly, suggesting that either high levels of relatedness or limited ability to discriminate the ancestral populations in the admixed-only SAMAFS data adversely affected the initial inference. Based on this augmented analysis, REAP and RelateAdmix have closer accuracies to that of PC-Relate yet remain somewhat less accurate (Supplemental Note; Supplemental Figure 4). The accuracy of PC-Relate and of KING are quite similar between the two analyses, with the exception that PC-Relate has improved accuracy for seventh degree relatives in the larger sample. Given this improvement and the fact that PC-Relate is the highest performing allele frequency-based method overall, we tested it further by varying its input parameters and the kinship values it uses to detect the set of individuals it uses to infer PCs. All these analyses resulted in similar accuracies except for different rates of inferred seventh degree relatives (Supplemental Note; Supplemental Figure 5); the variation in seventh degree relatedness inference may be due to stochastic factors and the relatively small numbers of these relatives in the dataset.

Besides considerations related to detecting population structure, the presence of many relatives in SAMAFS may lead to biased allele frequency estimates. Furthermore, haplotype phasing and therefore IBD inference accuracy might be greater than would be achieved in a sample composed mostly of unrelated individuals. To ensure the performance results presented here also apply to analyses of non-pedigree datasets, we identified a set of only distantly related individuals using FastIndep ^43^ and merged these samples with pairs of related individuals to form 1,000 datasets (Supplemental Note). Each reduced dataset contains at most one related pair of samples from any distinct SAMAFS pedigree, limiting the potential for bias. When classifying sample pairs included in at least one reduced dataset, PLINK’s inference accuracy differs by less than 3% for the first through fifth relatedness degrees compared to the full dataset (Supplemental Figure 2), suggesting that allele frequency biases are small and only minimally impact inference accuracy. In order to test the IBD detection methods, we increased the sample size of these reduced datasets by further merging 580 HapMap samples (Supplemental Note). Results from running the IBD segment-based methods on these datasets show a reduction in accuracy that ranges between 0% – 9.6% for first through fifth degree relatives, indicating that relatedness in SAMAFS may impact the inference accuracy (Supplemental Figure 3). Yet the results are still consistent with those of the larger analysis as the IBD segment-based methods generally have higher performance than allele frequency-based methods. This is true even in the reduced datasets that have no more than 1,204 samples and therefore are subject to a non-trivial rate of phasing error^44^.

In comparison to previous method evaluations, our results show some notable differences. For example, using real data from 30 pedigrees, ERSA reported lower accuracies for first through sixth degree relatives than we observe^19^, with differences ranging from 8.9% to nearly 21%. We believe this is attributable to differences in sample size, as the ERSA analysis considered only 304 individuals compared to 2,485 here. This—in addition to the accuracy reductions of IBD segment-based methods in the reduced datasets described above—indicates that sample size can have a dramatic impact on the quality of IBD segment-based methods. Thus smaller studies may wish to use allele frequency-based methods such as PC-Relate or, for non-admixed individuals, KING-robust, which in fact considers data from each sample pair separately rather than estimating allele frequencies from the full data ^17^. The authors of PC-Relate ^22^ find that KING and PLINK each tend to both overestimate and underestimate relatedness when analyzing admixed individuals, which is consistent with our results. They also report that PC-Relate generally outperforms REAP and RelateAdmix, matching our findings even after we incorporate additional HapMap individuals to aid detection of population structure (Supplemental Note). To our knowledge, other evaluations of relatedness inference have not included methods that directly detect IBD segments, and our results indicate that these are promising methods to apply in this setting.

As current methods provide only moderate accuracy when classifying third through seventh degree relatives, we evaluated the potential for increasing performance by combining inference results from the top three programs: ERSA, GERMLINE, and Refined IBD. We first used an approach that calls the degree of relatedness for a pair only when all three programs unanimously agree on the relatedness degree, providing no classification for other pairs (3,012 relative pairs and 632,615 reported unrelated pairs are unclassified). In comparison to the most accurate method’s performance in each degree class, the inference accuracy using this strategy increases only slightly for related pairs (+0.01%, +0.13%, +2.6%, +1.5%, +3.4%, +2.2%, and +1.1%, respectively, for first through seventh degree), but increases by 9.0% for unrelated pairs of individuals. This indicates a high level of discordance among the inferred relatedness status for a large fraction of pairs that are reported as unrelated. Many of these unrelated pairs must therefore have borderline inferences, and indeed most methods infer a sizeable fraction as only eighth degree relatives (Figure 1). We also considered a majority vote between the three programs, discarding cases in which all three programs inferred a different degree (only five relative pairs had such variable inferences while 110,848 pairs reported as unrelated are so discrepant). With this approach, there is a slight decrease in performance overall (-0.04%, -0.6%, -1.3%, -0.7%, -0.2%, -2.3%, and 0% for first through seventh degree relatives and +1.6% for unrelated samples). These results suggest that while there is room for improvement in the specificity of relatedness inference methods, dramatic improvement is likely to be achieved only with novel approaches and not composites of current methods. Of interest in this regard are recently developed methods that combine information across related individuals in order to infer a pedigree structure and/or improve relatedness accuracy^41,45,46^. Importantly, each of these methods relies on a pairwise relatedness approach, highlighting the continued relevance of pairwise inference methodologies even as new methods arise for addressing multi-way relatedness inference.

As an application of these findings, we leveraged the high accuracy of IBD segment-based methods to explore pairs of samples inferred to be closely related but reported as unrelated in the SAMAFS dataset. We used the top performing methods, ERSA, GERMLINE, and Refined IBD, to characterize unreported relatives. These three methods all infer a small number of first through third degree relationships that connect individuals from different pedigrees within SAMAFS (Figure 2; Supplemental Note). Overall, we found six pairs of pedigrees with at least five sample pairs between them that the methods unanimously infer to have first through third degree relatedness. Additionally, these three methods agree on the inference of 235 and 744 pairs of fourth and fifth degree relatives between the pedigrees (not shown), and suggest instances of reported first and second degree relatives likely to have the reverse relatedness class or to have much lower relatedness (Supplemental Table 3; Supplemental Note). These results highlight the necessity of checking reported or for unreported relatedness among samples in all cohorts and indicate that there can be sizeable numbers of unknown relatives across a range of relatedness degrees even in well-studied samples.

**Figure 2:**
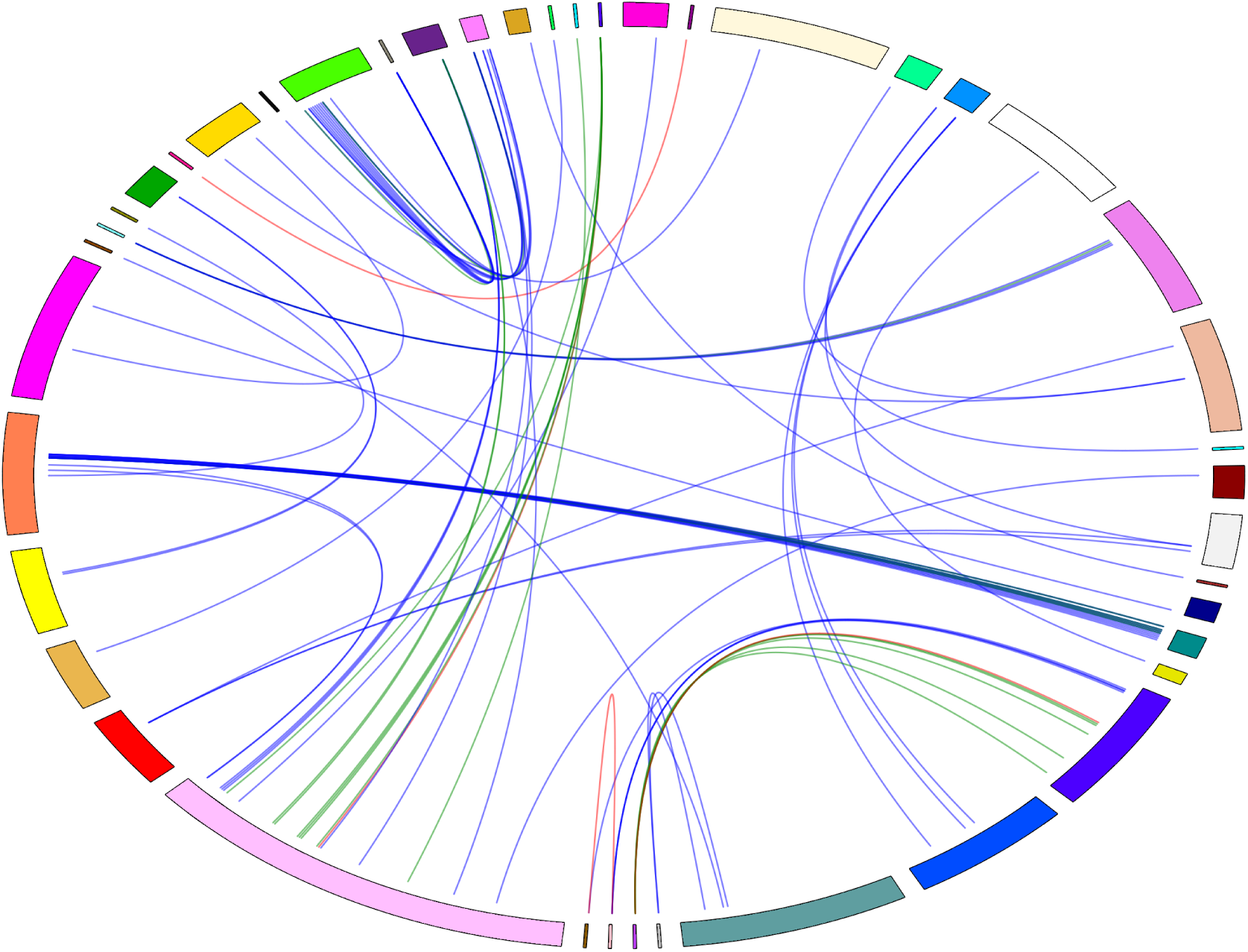
Relationships discovered between individuals from different SAMAFS pedigrees. Bands on the perimeter of the elliptical plot indicate distinct pedigrees within SAMAFS with band size proportional to the number of individuals in the pedigree. Curves between two bands correspond to discovered relative pairs with curve color indicating the degree of relatedness: red for first degree, green for second degree, and blue for third degree. Points where the curves end correspond to specific individuals, and a single point may have multiple curves running to it, indicating several relationships between that individual and others in the dataset.

Important factors for determining which analysis method to use in a study are its accuracy and its computational demands, and the runtimes of the methods evaluated here vary over several orders of magnitude (Table 2). PLINK is the fastest program with a runtime of only 18.1 seconds, while the IBD segment-based methods require up to 64 compute days in total (parallelized across 16 cores in our analyses). In general, we observe a trade-off between runtime and accuracy, with the top-performing methods being those that require the largest compute time, and with PLINK being one of the least accurate methods. Given the uniformly high accuracy of all methods for inferring first and second degree relatives, applications that are focused only on identifying close relatives have the option of using an efficient allele frequency-based method such as PLINK or PC-Relate to perform inference, the latter being an accurate program that is more computational intensive than PLINK but much faster than IBD segment-based methods. A further consideration is the ethnic group of the analysis cohort. PLINK and KING have biased results for distant relatives in the admixed SAMAFS data we focus on, but are expected to perform well in homogeneous populations or, for KING, collections of unadmixed samples from multiple homogeneous populations. On the other hand, for applications in which the aims include locating more distant relatives, the use of IBD segment-based methods should produce improved results. Although beyond the scope of this paper, recently developed methods for phasing extremely large samples ^47^ should improve upon the computational requirements of several methods (GERMLINE, ERSA, and HaploScore) and extend their utility to much larger datasets than the one we consider here.

We have presented a detailed comparison of state-of-the-art relatedness inference methods using thousands of pairs of individuals that range from first to seventh degree relatives as well as numerous sample pairs that are reported to be unrelated. All the methods we assessed reliably identify first and second degree relatives (accuracy ~92% – 99%), but their accuracy falls precipitously when classifying third to seventh degree relatives. This is unsurprising given the increased coefficient of variation as well as greater skewness in the proportion of genome shared as the meiotic distance between two relatives increases^15^. Despite these challenges, several IBD segment-based methods infer relatedness correct to within one degree of the reported relationship at a rate of ≥76.7% for all relationship degrees (Figure 1). Misreported or unknown relationships in the SAMAFS dataset likely explain some of the inference errors, particularly since even some confidently inferred first degree relationships were likely misreported as a more distant relationship or as unrelated (Supplemental Table 3; Figure 2). We find that IBD segment-based methods outperform other approaches for more distantly related pairs, though notably these packages require substantially more compute time to run (Table 2). While the precise performance results presented here are specific to the SAMAFS sample, we find that reducing the sample size still produces similar results, with methods that leverage IBD segments generally having greater accuracy than other approaches. Therefore, the results presented here should be generalizable to moderate and large scale studies and indicate overall properties of pairwise relationship inference methodologies: approaches that use IBD segments outperform other methods for third degree and more distant relatives; and the specificity of the inferences, even in a dataset where phase accuracy may be relatively high, are inhibited for all but the closest relatives.

## Data availability

The SAMAFS sample data are available on dbGaP under accession numbers phs000847 and phs001215.

## Acknowledgments

We thank the San Antonio Mexican American Family Study participants that made this analysis possible. We also thank Shai Carmi for helpful comments. This work was supported by a National Science Foundation Graduate Research Fellowship grant number DGE-1144153 to M.D.R.; Qatar National Research Fund grant NPRP 7-1425-3-370 to J.G.M.; an Alfred P. Sloan Research Fellowship, and a seed grant from Nancy and Peter Meinig to A.L.W.

